# Maternal bonding problems relate to aberrant neural processing of infant emotions during the first year postpartum: Results of an adapted fMRI Emotional GoNoGo Task

**DOI:** 10.1101/2024.11.30.625888

**Authors:** Monika Eckstein, Marlene Krauch, Ines Brenner, Beate Ditzen, Anna-Lena Zietlow

**Affiliations:** Institute for Medical Psychology, University Hospital Heidelberg, Heidelberg, Germany; Clinical Child and Adolescent Psychology, Faculty of Psychology, Technische Universität Dresden, Dresden, Germany

**Keywords:** imaging, cingulate cortex, emotion inhibition, bonding problems, postpartum depression, parental brain

## Abstract

Maternal bonding refers to the unique emotional connection between a mother and her baby that gradually develops during the peripartum period. However, 3-24% of women report bonding problems (BP), often accompanied by constraints for the mother-infant relationship, but not always depression with consequences for child development. Our present study investigates the neural and behavioral patterns that underlie the processing of emotional infant stimuli over the 1^st^ year postpartum parallel to a neurofeedback intervention. Mothers with and without BP (N = 45) completed a newly developed Emotional Infant GoNoGo Task while fMRI scanning at 3, 6 and 12 months postpartum. Our results show that response inhibition towards emotional infant faces elicits stronger results than towards adult faces in all mothers. While behavioral performance in BP is impaired, the neural responses to emotional infant faces as compared to neutral faces are increased at 3 months postpartum in limbic structures such as the anterior cingulate and insula, as well as nucleus caudatus. At 6 and 12 months behavioral reactions assimilate in BP to those of healthy controls, while differences in neural reactions between BP and healthy controls increase at 6 months and decrease again at 12 months. These effects are independent of depressive symptoms. Our findings point to an experience-based adaptive process of emotion processing and responses to infants’ affect during the first year postpartum as a specific characteristic of clinical BP. Implications are for potential therapeutic interventions to target emotion processing and regulation.

**Key Findings:**

- We present an adapted version of an Emotional GoNoGo Task for fMRI with infant stimuli: Behavioral inhibition towards emotional infant faces leads to increased activity in emotion regulation networks compared to adult faces in women during the 1^st^ year postpartum.
- Women with postpartum bonding problems (BP) show increased neural reactions (ACC, NCl Caudate, Insula) towards emotional baby faces (not adult faces, irrespectively of instructed inhibition and of valence) compared to a healthy control group (CG) at 3 months postpartum.
- On a behavioural level, women with BP show increased reaction times to positive emotional adult faces and higher error rates during the inhibition of reaction towards emotional baby faces at 3 months.
- Differences in neural reactions to emotional baby faces between BP and CG increase at 6 months and decrease again at 12 months following a neurofeedback intervention, while behavioral reactions in BD assimilate to those of CG

## Introduction

Maternal bonding refers to the unique emotional bond between a mother and her infant that already begins during pregnancy and continues to develop during the postpartum period (1, 2). Maternal bonding is evident behaviorally through cuddling, smiling at the infant, nurturing behaviors, and a high sensitivity to the infant’s needs (3). On a cognitive level, it involves the mother’s thoughts about the infant and his needs (4). On a neurobiological level, bonding is associated with alterations in structural and functional brain changes (5–7) and hormonal regulation (8). Strong maternal bonding can substantially improve infant development, influencing emotional and cognitive trajectories (for review see Le Bas, Youssef, (9)). Maternal bonding is related to neural sensitivity to infant-related stimuli, for example infant facial expressions (10, 11). The correct interpretation of infants’ facial cues is crucial because, compared to adults and older children, infants have limited social skills and primarily communicate through facial expressions, body language, and vocalisations (see Bowlby, (12, 13)). Therefore, correct and prompt interpretation of the infant’s expressions is essential for appropriate parental responses.

Research shows that healthy mothers demonstrate stronger and quicker neural responses to infant stimuli compared to non-mothers in crucial emotional processing regions such as the amygdala, insula, and orbitofrontal cortex. This response is particularly pronounced when interacting with their own infants (14, 15). This increased responsiveness to infant cues can be partly explained by neurobiological changes during transition to parenthood (6, 16, 17) and also includes an altered response to infant stress cues (18). Functional and structural changes are observed in core limbic and cortical socio-cognitive networks that are unique to parents and related to caregiving behavior (17, 19, 20). It is assumed that these alterations endure (see Orchard, Rutherford (21) for review).

Healthy mothers with a close maternal-infant bond show specific patterns of neural activity when responding to negative infant stimuli. In particular regions such as the amygdala, anterior cingulate cortex (ACC), ventral prefrontal cortex (PFC) and insular cortex are associated with emotional processing and regulation of negative infant stimuli (15, 22). On a neural level, a child’s face triggers stronger activation in the fusiform gyri, the middle occipital gyri, the superior temporal gyri, the supplementary motor area, the pre- and postcentral gyri, as well as in the amygdala, when compared to an adult’s face (23). However, previous studies differ in the methods they use (own or unfamiliar, visual or acoustic, infant or adult facial stimuli). E.g., Rupp et al (24) investigated mothers compared to non-mothers when exposed to negative emotional infant images and report reduced activation in the amygdala and reduced subjective negative arousal in mothers. Similarly, in an Emotional GoNoGo Task, mothers showed increased NoGo P3 amplitudes compared to non-mothers, underlining differences in emotional regulatory mechanisms between mothers and non-mothers (18).

Research in this area underscores the significance of emotion regulation in tempering over-reactivity to infant’s crying, a factor closely associated with maternal mental health (25, 26). Emotion regulation can be considered a higher-order process that follows the initial steps of direct emotion processing (i.e., attention and perception of information eliciting emotional arousal) and the subsequent reaction to these stimuli. Inhibition of a direct response in order to avoid negative consequences requires medium-level brain circuitries, such as the limbic system including the cingulate cortex, but not cognitive executive function (27). Proficiency in utilizing multiple strategies to regulate emotions may serve to prevent high parental stress and mental illness (28). Specifically mothers’ sensitivity to their infants’ facial expressions is correlated with activation in brain regions associated with emotion regulation, particularly behavioral inhibition (18), while emotion regulation in a broader sense seems also impaired in postpartum mental disorders such as depression and anxiety (29, 30).

Maternal characteristics and health are thought to affect the neural processing of infant stimuli, e.g. maternal sensitivity influences neural activation in response to infant stimuli ((31–35)). Mothers who were more sensitive in the interaction with their child, showed greater activation in the right frontal pole and inferior frontal gyrus when listening to their own baby crying as compared to the unfamiliar infant 18 months postpartum. Mothers interacting less sensitive showed higher activation in the left insula and temporal pole [35]. Regarding postpartum depression several studies suggest a reduced processing of infant faces (36–38). For instance, Moses-Kolko et al., (38) investigated neural reactions to negative facial expressions using fMRI in both mothers diagnosed with postpartum depression and a control group of healthy individuals. The results suggest that reduced activity in the dorsomedial PFC and reduced effective connectivity between the dorsomedial PFC and the amygdala in response to negative emotional faces may reveal a crucial neural mechanism or consequence of postpartum depression.

The presence of a maternal BP may be another possible influential factor for impaired infant emotion processing. A notable portion of women, specifically 3 to 24 % in community samples (3), (39–45) face challenges in forming a good emotional bond with their offspring. While BD can indeed manifest in healthy mothers, it occurs more frequently in the context of perinatal mental illness such as perinatal depression, anxiety, or post-traumatic stress disorder (42, 46, 47). For mothers with perinatal depression, the prevalence rates of BD range from 17-29% (48). BD shows in various dimensions, including emotional disconnection, and feelings of rejection or frustration towards the infant (43, 49, 50). As maternal caregiving behavior may be restricted in mothers with BP (51) this is probably paralleled by neural alterations. Accordingly, mothers who reported poorer postpartal bonding have been reported to process stressed infant faces slower in late pregnancy (52). In addition, faster processing of infant faces compared to adult faces in an EEG task during pregnancy has been reported to be associated with higher maternal bonding quality (11). Behavioral inhibition towards social emotional stimuli can be assessed using a GoNoGo paradigm that requires the individual to correctly identify emotions and flexibly change his/her reaction to them. As task performance is often impaired in pathological samples (53, 54), a non-social GoNoGo Task is an established fMRI paradigm. The extension to adult facial stimuli is utilized especially in disorders that are characterised by social deficits (e.g. depression (55)).

However, BP in particular has so far not yet been systematically investigated in terms of behavioral and neural responses to emotional child-related stimuli. Yet, understanding the processing of emotional infant stimuli and subsequent maternal emotion regulation may help to explain the parenting difficulties of mothers with BP. Based on the state of research summarized above, we expect behavioral and neural deficits processing infant and adult faces within a GoNoGo Task in mothers with BP compared to a control group.

The present study is part of a larger project including a fMRI-neurofeedback intervention (56). Details and results of the intervention part are prepared for publication elsewhere. Here we present results of an adapted version of the Emotional GoNoGo Task for fMRI using infant stimuli. We investigate the neural mechanisms underlying inhibitory responses to emotional infant faces compared to adult faces in women at 3, 6 and 12 months postpartum. We analyze behavioral outcomes associated with BP, focusing on reaction times and error rates during inhibition of responses to emotional baby faces compared to emotional adult faces in order to describe the neurobehavioral response patterns in women with BD to emotional infant stimuli during the postpartum period and disentangle them from depressivity. Based on the state of research summarized above, we expect to detect both, group differences and differences for the progress of the postpartum period, in behavior and neural key nodes related to parental emotion regulation such as the ACC.

## Methods

N = 64 participants were recruited as part of a larger study on postpartum bonding disorders (for details see 56, approval was given by Ethics Committee of the Medical Faculty Heidelberg, S-450/2017). Recruitment was done mainly via data of registration offices. Following initial contact of 461 mothers, a phone-screening interview with 254 was administered 4 to 10 weeks postpartum to address preliminary exclusion criteria (see Figure 1). Exclusion criteria were severe psychiatric conditions that needed treatment, substance abuse, pre-term birth, multiple birth, severe health issues of the baby and MRI contraindications.

**Figure 1.**
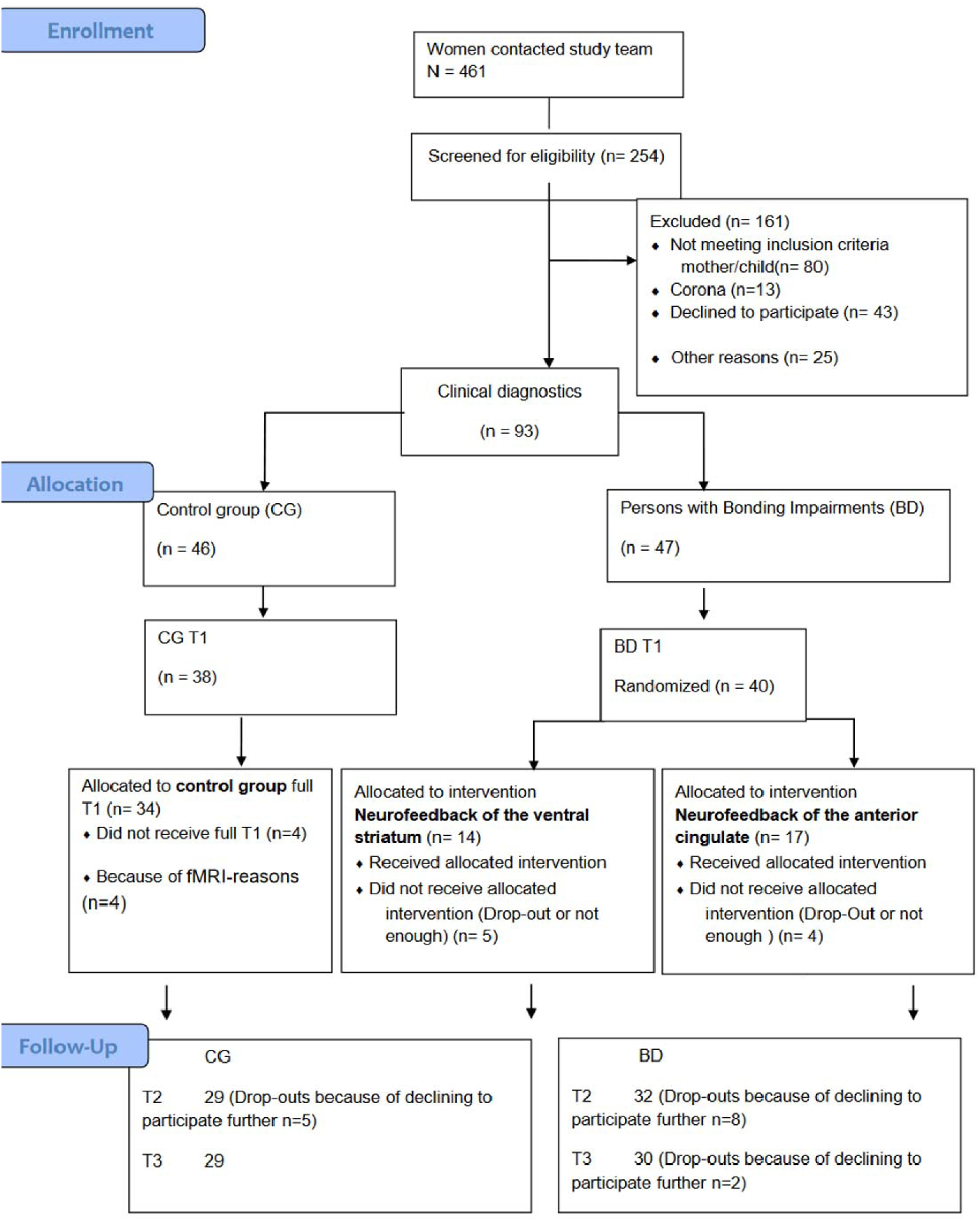
Project Participant Flowchart for

Subsequently, based on the results of the mother’s bonding interview conducted with 93 based on the proposed criteria at T1, mothers were assigned to either the intervention (BP, n=40) or control group (CG, N = 34). All participants were subjected also to detailed psychodiagnostics according to DSM-5 over the course of the study. Participants performed three task-based fMRI sessions over the first year postpartum: with approx.3 months (T1), 6 months (T2) and 12 months (T3) postpartum. During these sessions, they first performed a monetary reward task, subsequently the Infant Emotional GoNoGo Task and a passive viewing task of pictures of their own child and partner. The BP group received a neurofeedback intervention between T1 and T2. Results of the intervention and other tasks will be published separately.

### fMRI task: Infant Emotional GoNoGo Task

Using a specifically adapted GoNoGo paradigm, participants were presented with positive, negative and neutral expressions of pictures of unknown babies, aged approx. 4-10 months, and, unknown adults with positive, negative and neutral expressions, as well as non-social control stimuli (geometric figures; a circle, a cross, a diamond and a triangle) over six presentation blocks. Facial stimuli were taken from an established database (57) with positive, neutral and negative affect in infants with age 3-5 months and the Karolinska faces for adults, see Figure 2.

**Figure 2.**
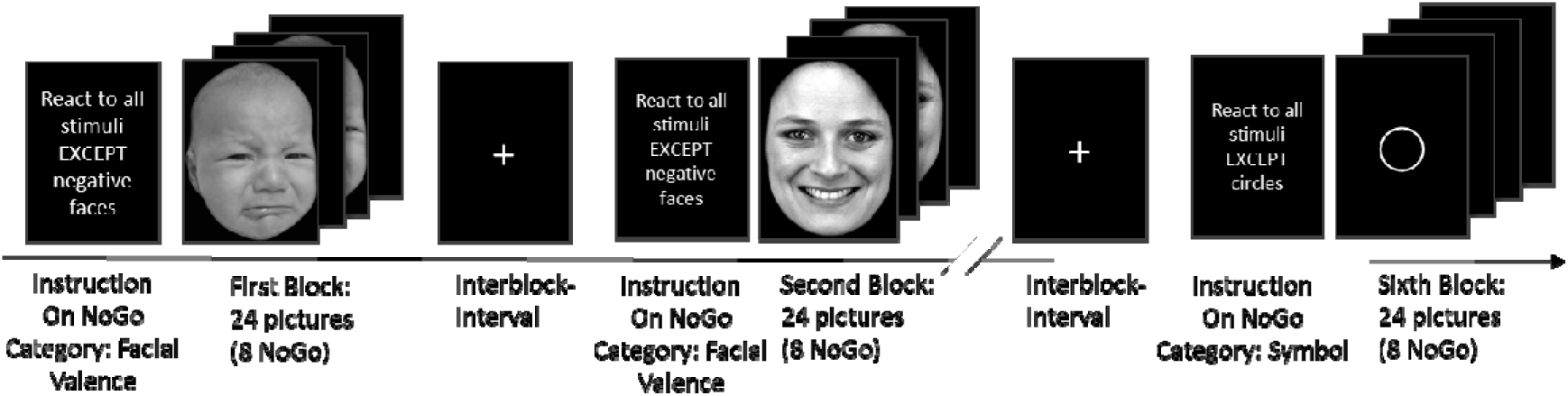
fMRI paradigm: the Infant Emotional GoNoGo Task comprising six blocks in randomized order (stimuli from available data bases 57).

The following factors were systematically manipulated: child versus adult and emotionality of facial expression (positive vs negative vs. neutral). In two blocks, the participants received instructions to respond by pressing a button as fast as possible (Go trials) to all facial expressions except (NoGo trials) the negative (one block babies, one block adults). In two other blocks, they were instructed to respond as fast as possible to all except the positive stimuli (one block babies, one block adults). In the two non-social blocks, the participants were instructed to react as fast as possible to all shapes but not to a circle or a diamond.

Each block consisted of 12 pictures shown twice for 500ms, therefore 24 trials, of which 8 were NoGo trials. Fixation cross between trials was jittered from 1500-2000ms. Between blocks was an interval of 5000ms. Total task duration was approx. 14 minutes.

### Self reports

The assessment of bonding problems was conducted using an interview based on the criteria outlined by Brockington (48). These criteria categorize bonding problems into Delay/Loss of Bonding, Pathological Anger (mild/moderate/severe) and Rejection (threatened/established), which may present with additional symptoms when combined. Maternal depression was screened using the Edinburgh Postnatal Depression Scale (EPDS) (58).

### fMRI data acquisition

Imaging was performed using a 3Tesla Prisma-Fit Siemens Scanner (Siemens, Erlangen, Germany) at the Department of Neuroradiology at University Hospital Heidelberg. First a detailed anatomical scan was obtained with a magnetization prepared rapid gradient echo (MPRAGE) sequence with repetition time TR=1.9s, echo time TE=2.52ms, flip angle=9° and an isotropic resolution of 1x1x1mm, followed by functional scans. Functional images were acquired with an Echo Planar Imaging (EPI) sequence with TR=1.64s, TE=30ms, flip angle=73° and GRAPPA factor 2 in 30 slices of 3mm thickness, and field of view FoV=192mm for a voxel size of 3x3x3mm.

### Analyses

Behavioral data were analysed using IBM SPSS statistics 29.0 using t-tests for independent groups (BP vs. CG) with a two-tailed p<.05. We additionally performed one-way analyses of variance with depressiveness (EPDS) as a covariate (ANCOVAs) to control for depression effects on group differences.

For fMRI data, using SPM12 (Wellcome Center for Human Neuroimaging, London, UK) we first conducted preprocessing of the functional data with the following steps: Slice time correction, realignment to the first image. Anatomical images were segmented and normalized to the SPM 12 NMI template. Functional images were coregistered with these anatomical images and normalized in NMI space rescaling to voxel size 2x2x2mm. After smoothing with a full width at half maximum FWHM=8x8x8mm Gaussian kernel, additional normalization to NMI space with rescaling to 1x1x1mm was done. The first 5 images of each session were discarded.

On the first level, we specified an event-related model with 3 sessions and for each the 12 task conditions as saved in the logfiles with the onsets of symbols, adult positive faces, adult negative faces, baby positive faces and baby negative faces, both as Go and NoGo conditions, and neutral adult and neutral baby faces as Go condition. Interstimulus intervals entered the implicit baseline. We controlled for the movement regressors obtained from preprocessing and applied a high pass filter of 128 and convolved the conditions with the canonical hemodynamic response function. The estimated model calculated contrasts for the relevant conditions such as T1[emotional baby face > neutral baby face] and Mean[[baby face NoGo > baby face Go] > [adult face NoGo > adult face Go]]. On the second level, participant-specific contrast maps from the first level analyses were compared between patients and controls with two-sample t-tests and EPDS scores were tested as covariates. A significance threshold of p=0.05 cluster-level FWE-corrected with cluster-defining height threshold of p=0.001 uncorrected was applied. Probabilistic labelling of regions for the tables was done using SPM built-in Neuromorphometrics atlas.

## Results

### Sample characteristics

N = 64 started in the overall project. Due to the Covid-19 pandemic, several fMRI sessions for T2 and T3 had to be cancelled, resulting in diverging sub-samples for some of the analyses.

**Table 1.** Sample description for participants that completed T1.

|  |  | Mothers with bonding problems (BP) |  |  | Control group (CG) |  |  | Difference |
| --- | --- | --- | --- | --- | --- | --- | --- | --- |
|  |  | N | Mean | SD | N | Mean | SD |  |
| Depressiveness (EPDS) | T1 | 31 | 10.068 | 4.53 | 28 | 3.036 | 2.78 | T(57)=7.11, p = 0.021* |
|  | T2 | 24 | 7.417 | 4.95 | 23 | 2.522 | 2.73 | T(45)=4.32, p <0.001* |
|  | T3 | 23 | 5.043 | 3.78 | 25 | 3.64 | 2.75 | T(46)= 1.48, p = .073 |
| Bonding Problems (PBQ) | T1 | 31 | 24.709 | 11.21 | 28 | 5.607 | 3.39 | T(57)=8.66, p <0.001* |
|  | T2 | 26 | 15.077 | 6.09 | 26 | 5.039 | 2.27 | T(50)=7.54, p < 0.001* |
|  | T3 | 23 | 15.043 | 9.65 | 23 | 6.68 | 5.52 | T(46)=3.72, p <0.001 |
| Maternal age at delivery |  | 30 | 32.93 | 3.54 | 28 | 31.39 | 3.45 | T(56)=1.67, p = 0,869 |

Abbrev: PBQ, Postpartum Bonding Questionnaire (Reck et al, 2006), \* significant result

N= 55 completed the first fMRI Session (T1), N=31 mother with BP, N=28 healthy controls. Complete data of N = 45 persons for the three fMRI sessions at 3, 6 and 12 months was available that entered the analyses reported below. Of these 45 mothers, N = 19 built the healthy control group and N = 26 the patient group with bonding problems. The patient group was further allocated to either an index neurofeedback intervention or a control neurofeedback intervention between 3 and 6 months (as described in 56). No neural or behavioural differences between these two intervention groups were observed for the present fMRI task. Therefore data was pooled. Focus of interpretation is the results of T1 that are not influenced by the intervention.

In the BP group, 12 mothers (36.36 %) reported current depression, 5 mothers (15.15%) were diagnosed with a current anxiety disorder, and 9 mothers (27.27 %) had depression with comorbid anxiety. For lifetime diagnoses, 2 mothers (6.06 %) reported lifetime depression, and in each case, one mother had an anxiety disorder or depression with comorbid anxiety disorder. From the BP group, 3 mothers (9.09 %) did neither meet criteria for any current nor lifetime mental disorder.

### Behavioral results

In the emotional Go condition, the t-test for paired samples over all participants, mothers with BP and healthy participants, revealed no significant difference in reaction times to emotional adult faces vs. emotional baby faces.

Regarding group differences between mothers with BP and healthy participants, an independent samples t-test revealed significantly longer reaction times towards positive emotional adult faces in mothers with BD compared to CG (T(53) = 2.96, p = 0.002, Figure 3). This group difference remained significant after adjusting for depressiveness, measured with EPDS (F(1,48) = 7.83, p = 0.007, partial η² = 0.140). There were no significant differences between groups for negative emotional adult faces, emotional baby faces or non-social control stimuli regarding reaction times.

**Figure 3.**
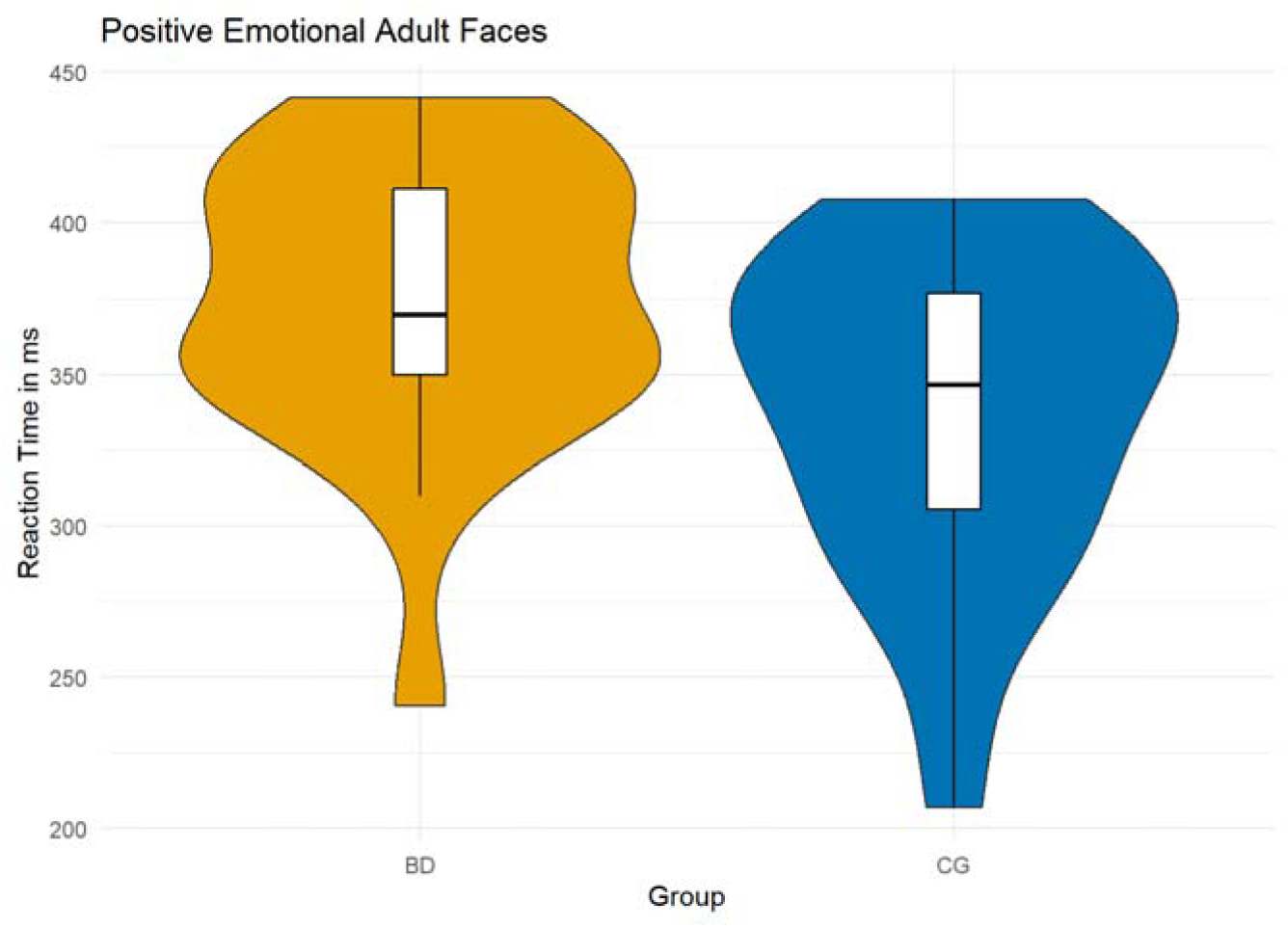
Go condition: Mean reaction times for positive emotional adult faces at T1, mothers with PD (MEAN = 376.2 SD = 44.37) vs. healthy controls (MEAN = 338.7, SD = 49.61).

In the emotional NoGo condition, the t-test for paired samples over all participants revealed significantly higher error rates for emotional baby faces compared to emotional adult faces (T(44) = -3.4, p = 0.001).

Regarding group differences between mothers with BP and healthy participants, the t-test revealed significantly higher error rates for emotional baby faces in mothers with BP compared to healthy controls (T(50) = 2.0, p = 0.026, Figure 4). This group difference remained significant after adjusting for depressiveness, measured with EPDS (F(1,46) = 5.51, p = 0.023, partial η² = 0.107). There were no significant differences between groups for adult emotional faces or non-social control stimuli regarding error rates.

**Figure 4.**
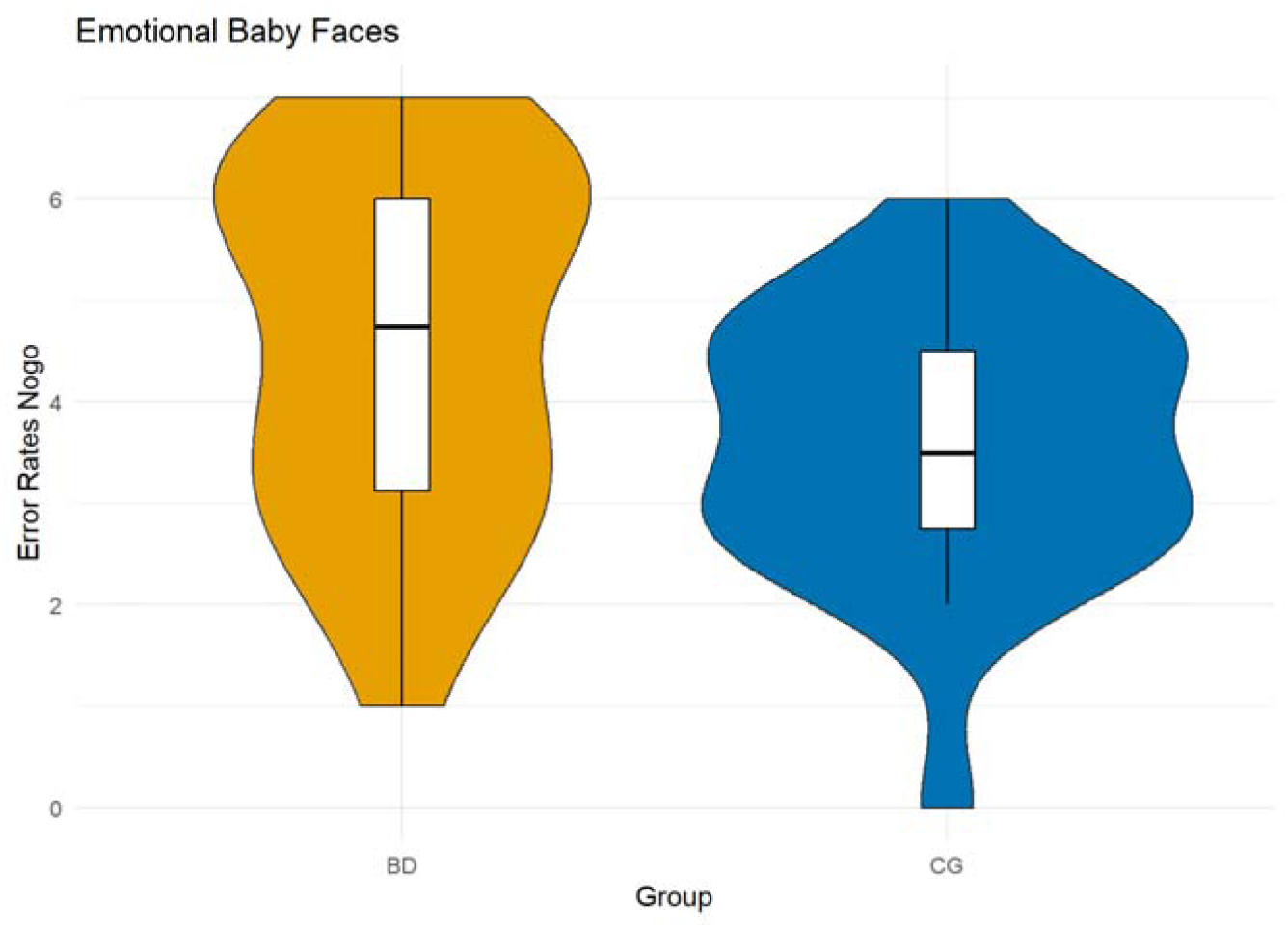
NoGo condition: Mean error rates for emotional baby faces at T1, mothers with BD (MEAN = 4.8, SD = 1.78) vs. CG (MEAN = 3.9, SD = 1.58).

At 6 and 12 months, groups did not differ significantly for emotional baby or adult facial stimuli or non-social control stimuli regarding reaction times or error rates.

### fMRI results

All analyses were first conducted with EPDS scores as covariate, which did not reach any significance. Therefore, we report the parsimonious results without EPDS.

### Main effects of task

The Infant Emotional GoNoGo Task activated a broad network of emotion processing and emotion inhibition over all participants and time points as assessed by contrasting [baby faces NoGo > baby faces Go] irrespectively of emotional valence. Among others, significant activations were found in anterior and middle cingulate cortex, inferior frontal gyrus, temporal gyrus and insula, see Figure 5 and Table 2.

**Figure 5.**
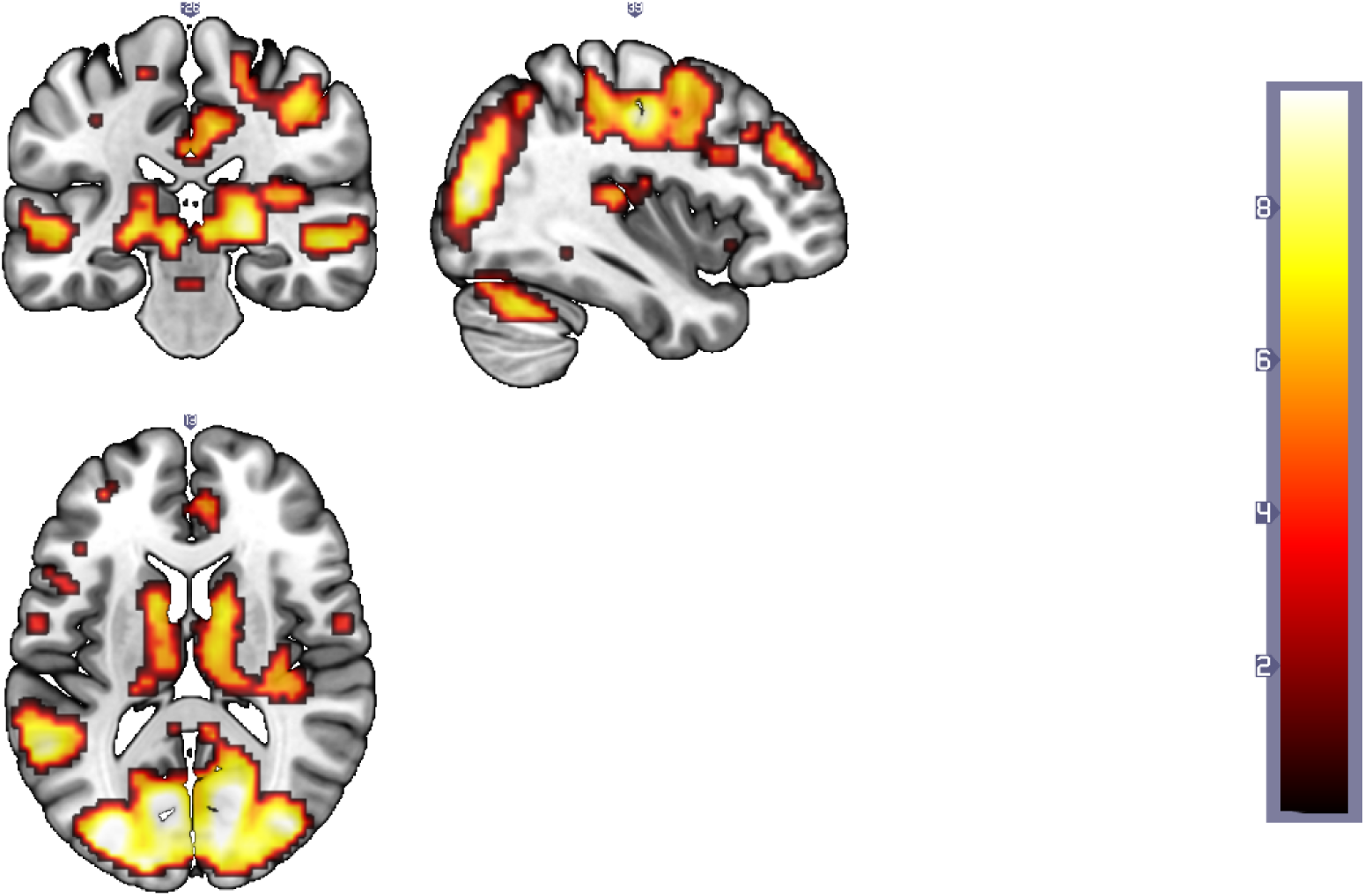
Neural activation of inhibition towards NoGo baby stimuli as compared to non-inhibition (=reaction, Go) over all participants and assessments

**Table 2.** Activations pattern for the baby NoGo condition as compared to the Go condition.

| cluster | cluster size | peak | peak | peak | NMI coordinates (mm) |  |  | hemisphere | region |
| --- | --- | --- | --- | --- | --- | --- | --- | --- | --- |
| p(FWE-corr) | equivk | p(FWE-corr) | T | equivZ | x | y | z |  |  |
| 0.00 | 10231.00 | 0.00 | 15.81 | 65535.00 | -15 | -91 | -4 | L | calcarine cortex |
|  |  | 0.00 | 15.26 | 65535.00 | 18 | -91 | -1 | R | calcarine cortex |
|  |  | 0.00 | 15.21 | 65535.00 | -9 | -85 | 2 | L | calcarine cortex |
| 0.00 | 758.00 | 0.00 | 12.58 | 65535.00 | -42 | 2 | 47 | L | precentral gyrus & middle frontal gyrus |
|  |  | 0.00 | 11.14 | 7.56 | -45 | -4 | 35 | L | precentral & postcentral gyrus & middle frontal gyrus |
|  |  | 0.00 | 8.62 | 6.51 | -48 | 14 | 23 | L | inferior frontal, middle frontal and precentral gyrus |
| 0.00 | 673.00 | 0.00 | 9.74 | 7.01 | -3 | 8 | 62 | L | supplementary motor cortex and superior frontal gyrus |
|  |  | 0.00 | 7.23 | 5.79 | 9 | 14 | 56 | R | supplementary motor cortex and superior frontal gyrus |
|  |  | 0.00 | 7.18 | 5.77 | 6 | 26 | 35 | R | supplementary motor cortex<br>anterior and middle cingulate and superior frontal gyrus |
| 0.00 | 147.00 | 0.00 | 8.29 | 6.35 | 54 | -25 | -7 | R | middle and superior temporal gyrus |
|  |  | 0.00 | 7.33 | 5.85 | 63 | -28 | -4 | R | middle and superior temporal gyrus |
| 0.00 | 43.00 | 0.00 | 6.94 | 5.63 | 54 | -46 | 26 | R | angular and supramarginal gyrus |
| 0.00 | 42.00 | 0.00 | 6.82 | 5.57 | -30 | 35 | 35 | L | middle and superior frontal gyrus |
| 0.00 | 15.00 | 0.00 | 6.65 | 5.47 | -42 | 44 | 20 | L | middle frontal gyrus |
| 0.00 | 49.00 | 0.00 | 6.56 | 5.42 | 54 | -4 | -10 | R | superior and middle temporal gyrus |
|  |  | 0.02 | 5.49 | 4.74 | 60 | 5 | -4 | R | superior temporal gyrus and temporal pole |
| 0.00 | 22.00 | 0.00 | 6.11 | 5.14 | -54 | -7 | -10 | L | superior and middle temporal gyrus |
| 0.00 | 17.00 | 0.01 | 6.01 | 5.08 | 9 | -13 | 53 | R | supplementary motor area and precentral gyrus |
| 0.01 | 6.00 | 0.01 | 5.92 | 5.02 | 0 | -37 | -31 |  | brain stem |
| 0.02 | 2.00 | 0.01 | 5.64 | 4.84 | -24 | 8 | -13 | L | putamen and anterior insula |
| 0.00 | 13.00 | 0.02 | 5.58 | 4.80 | -18 | -37 | 59 | L | postcentral gyrus and superior temporal lobule |
| 0.01 | 3.00 | 0.02 | 5.57 | 4.79 | -30 | 50 | 11 | L | middle and superior frontal gyrus |
| 0.01 | 5.00 | 0.02 | 5.56 | 4.78 | 54 | 26 | 26 | R | middle and inferior frontal gyrus |
| 0.01 | 4.00 | 0.02 | 5.54 | 4.77 | 54 | -49 | -10 | R | inferior and middle frontal gyrus |
| 0.01 | 3.00 | 0.03 | 5.43 | 4.70 | 54 | 14 | -7 | R | temporal pole and frontal operculum |
| 0.01 | 3.00 | 0.03 | 5.42 | 4.69 | 57 | 23 | 20 | R | inferior and middle frontal gyrus |
| 0.01 | 4.00 | 0.03 | 5.41 | 4.68 | 0 | -25 | -22 |  | brain stem |
| 0.01 | 3.00 | 0.03 | 5.36 | 4.65 | -18 | -25 | 59 | L | precentral and postcentral gyrus |
| 0.03 | 1.00 | 0.03 | 5.35 | 4.64 | 36 | 20 | -7 | R | anterior insula and inferior frontal gyrus |
| 0.03 | 1.00 | 0.04 | 5.30 | 4.61 | 9 | -4 | 47 | R | supplementary motor area and middle cingulate cortex |
| 0.03 | 1.00 | 0.04 | 5.28 | 4.59 | -9 | -10 | 38 | L | middle cingulate cortex and supplementary motor area |
| 0.02 | 2.00 | 0.04 | 5.23 | 4.56 | 54 | -43 | 41 | L | supramarginal and angular gyrus |

Comparing reactions toward adult faces with reactions towards baby faces [baby faces NoGo > baby faces Go] > [adult faces NoGo > adult faces Go] resulted in even stronger activation patterns for the baby stimuli in posterior cingulate, hippocampus and other key nodes for emotion processing, see Figure 6 and Table 3.

**Figure 6.**
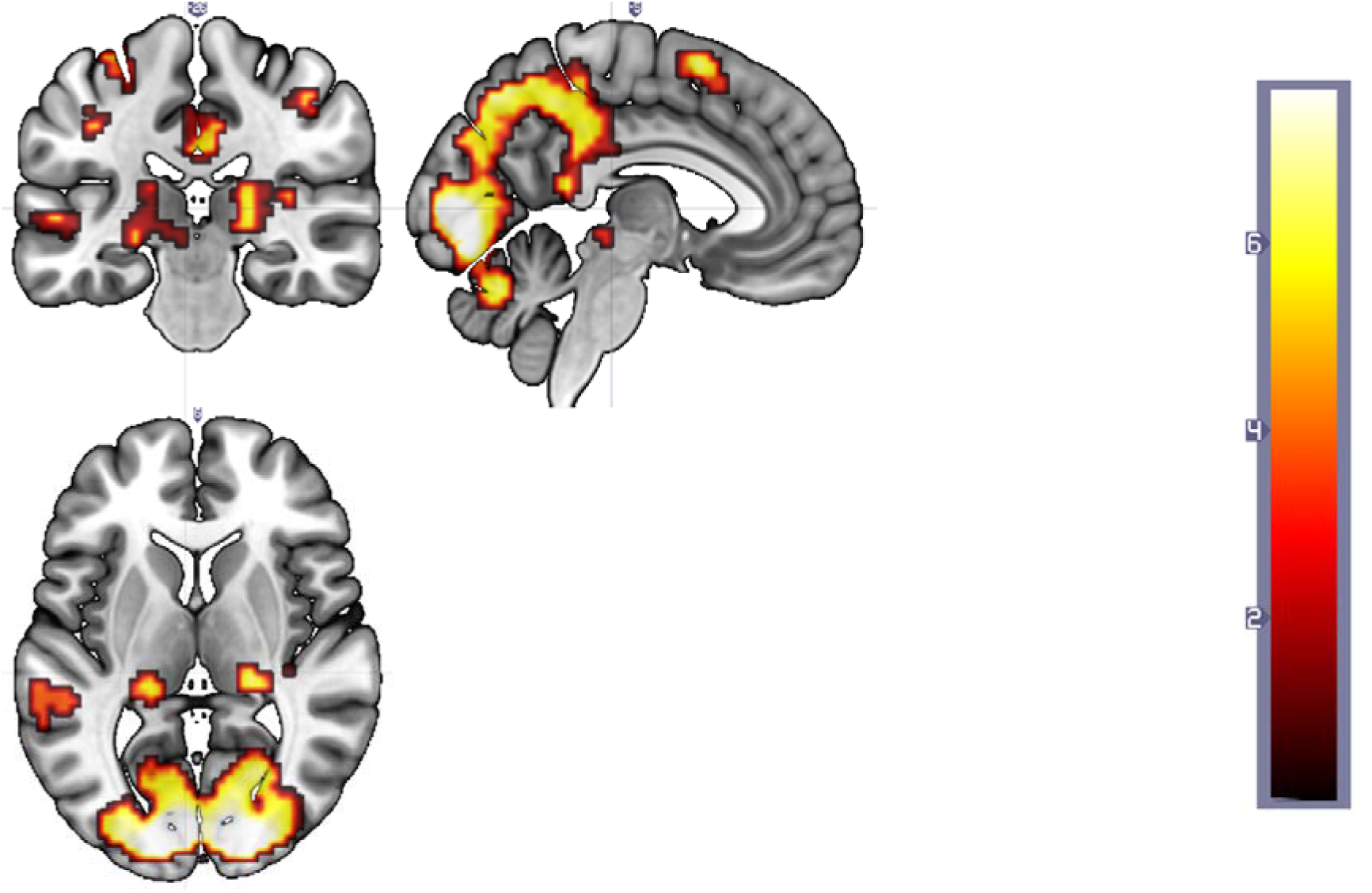
Neural activation of (inhibition towards baby stimuli as compared to non-inhibition) as compared to (inhibition towards adult stimuli as compared to non-inhibition) over all participants and assessments.

**Table 3.**
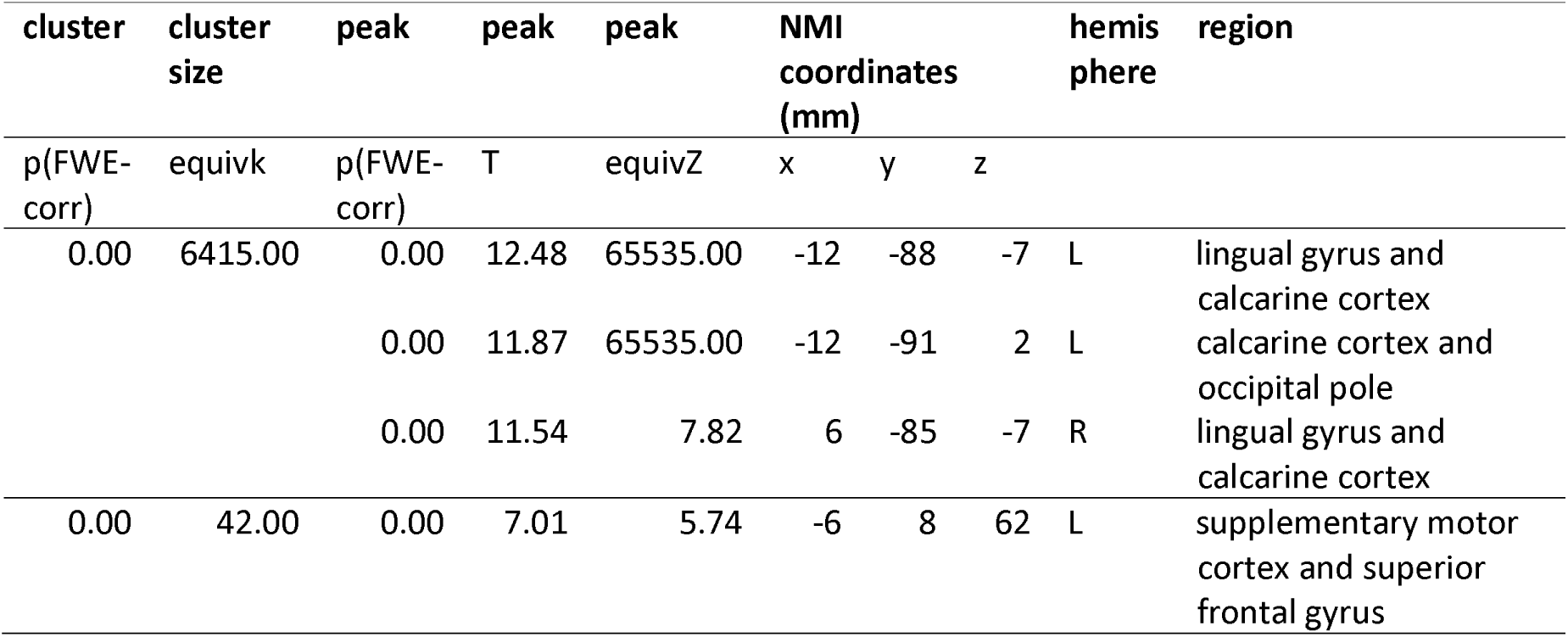
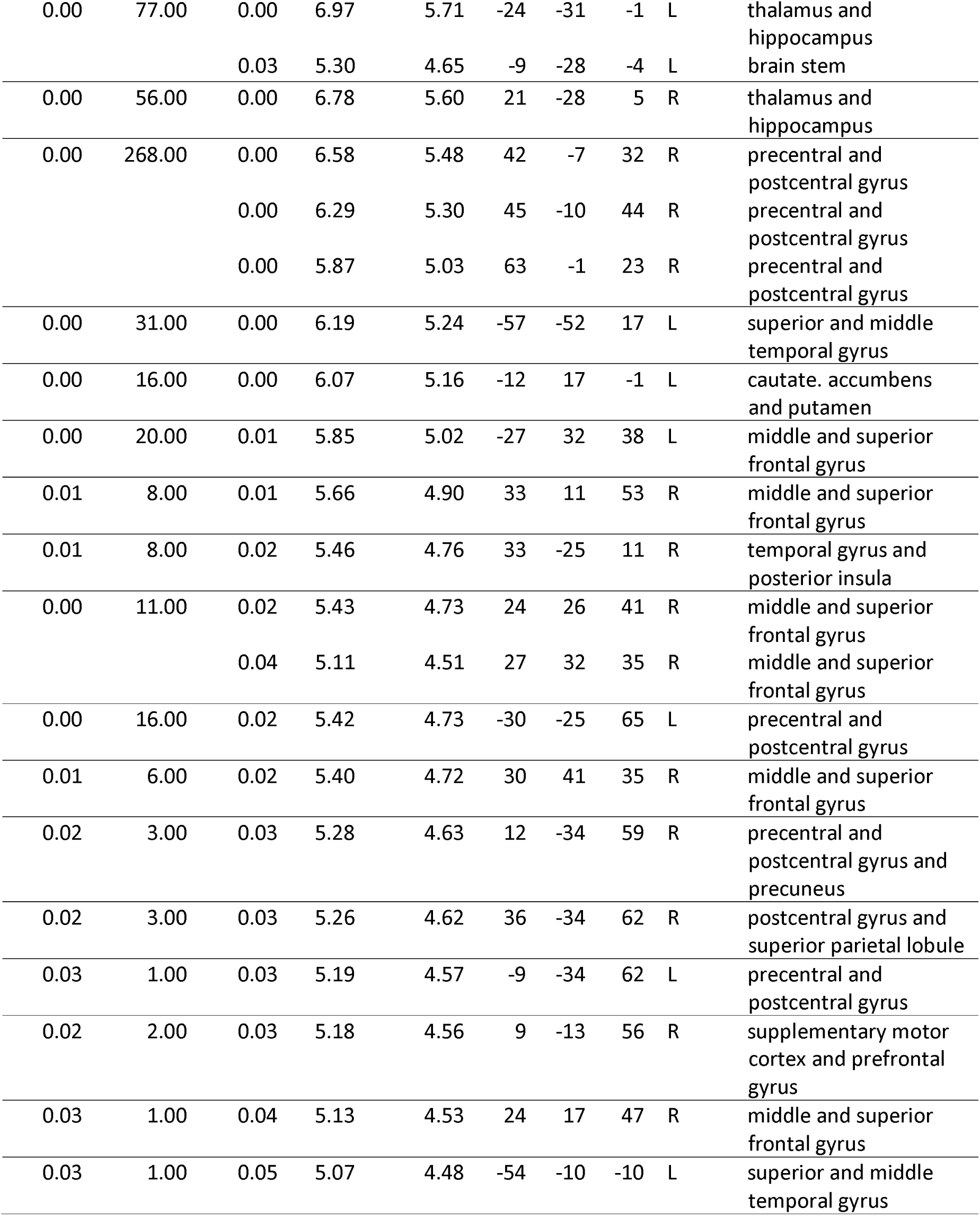
Activation pattern for the baby NoGo condition as compared to the Go condition.

### Group comparison

#### 3 months postpartum

Comparing the groups at their first assessment (T1) shows a significant difference in processing emotional baby faces, irrespectively of Go or NoGo instruction and of valence. BP participants showed stronger reactions in midline areas such as anterior and medial cingulate and basal ganglia of the dorsal striatum, see Figure 7 and Table 4. No group differences or interactions gain significance for adult faces or symbols.

**Figure 7.**
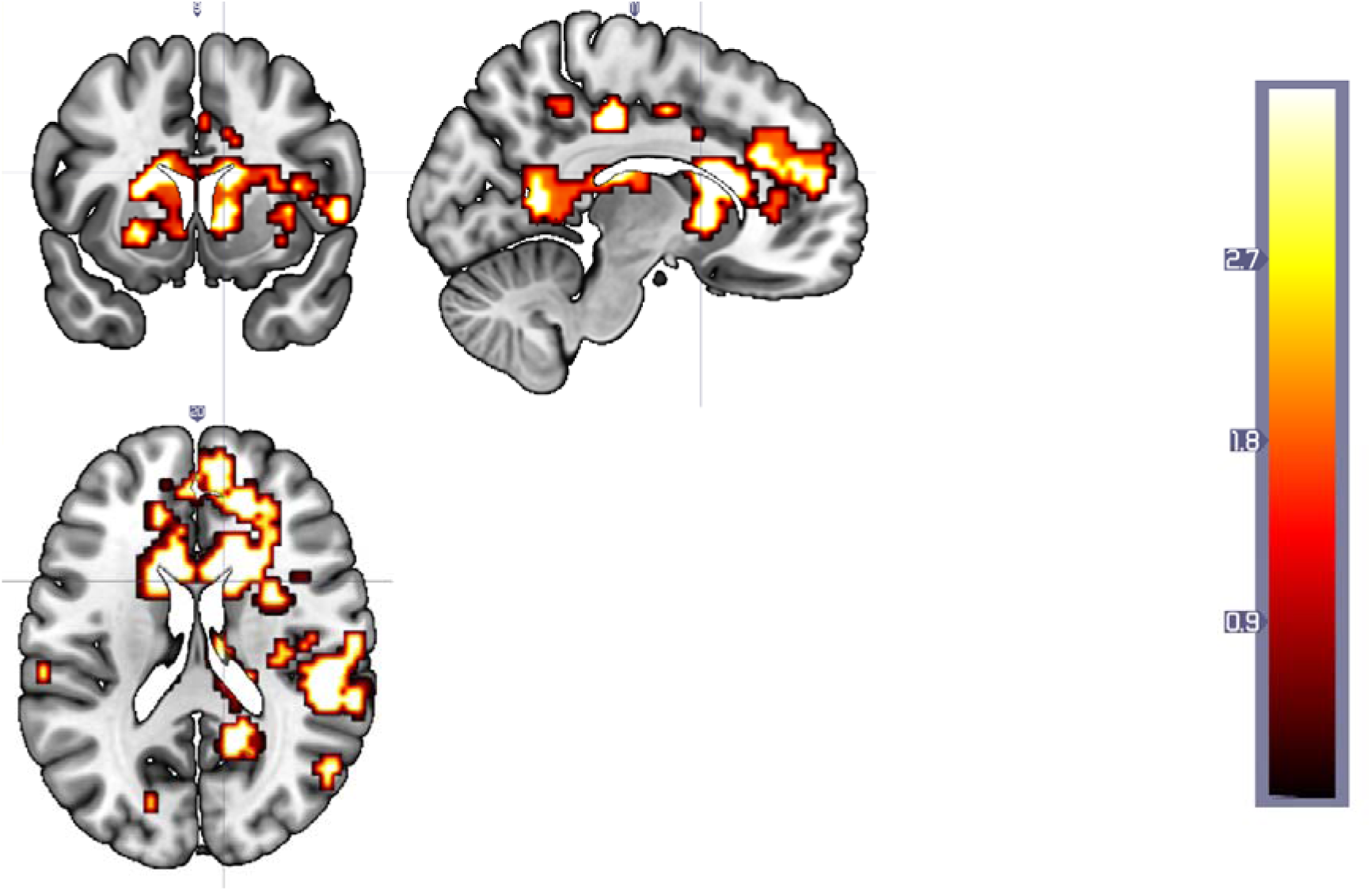
Increased neural activation at T1 in patients (BP) while processing emotional baby stimuli as compared to neutral baby stimuli as compared to healthy controls.

**Table 4.**
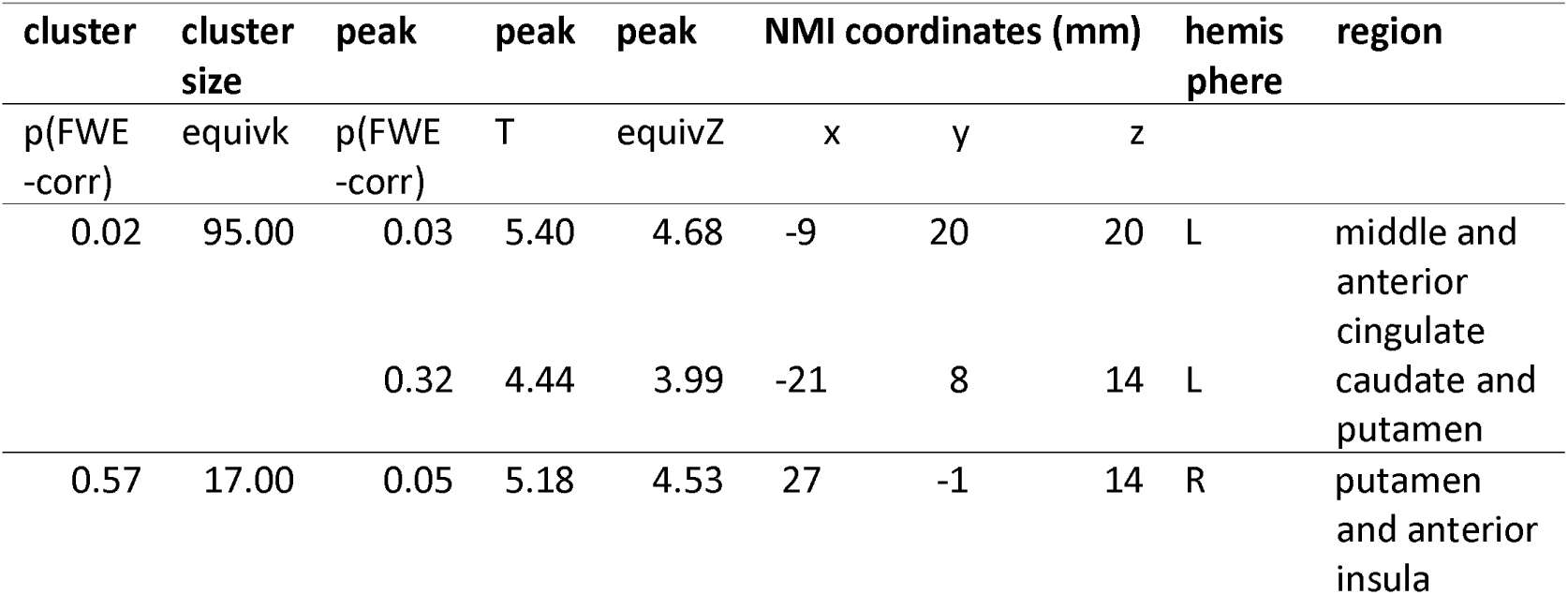
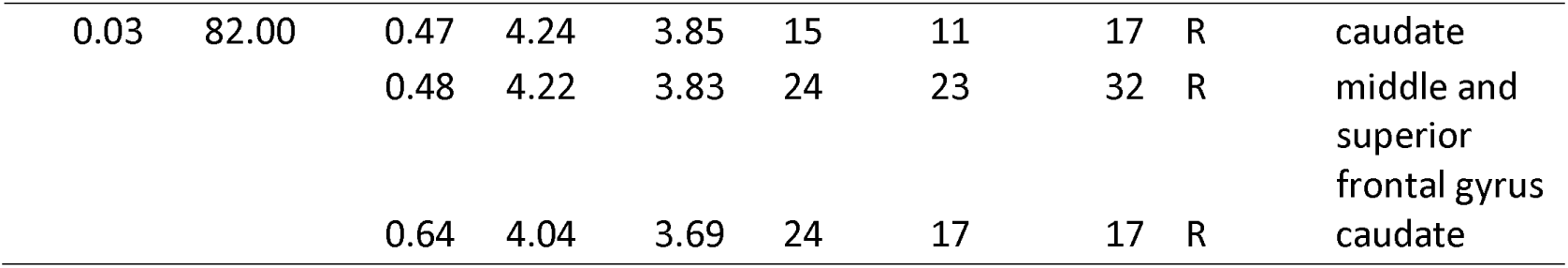
Activations pattern for BP larger than CG in the response to emotional baby faces as compared to neutral baby faces at 3 months postpartum.

#### 6 months postpartum

Comparing the differences of the first (T1) and second (T2) assessment between groups there is even an increase in the difference. While processing emotional baby faces, irrespectively of Go or NoGo instruction and of valence, BP mothers increase their response in frontal gyruses, cingulate, caudate and temporal areas, see Figure 8 and Table 5. They had received an intervention between these assessments.

**Figure 8.**
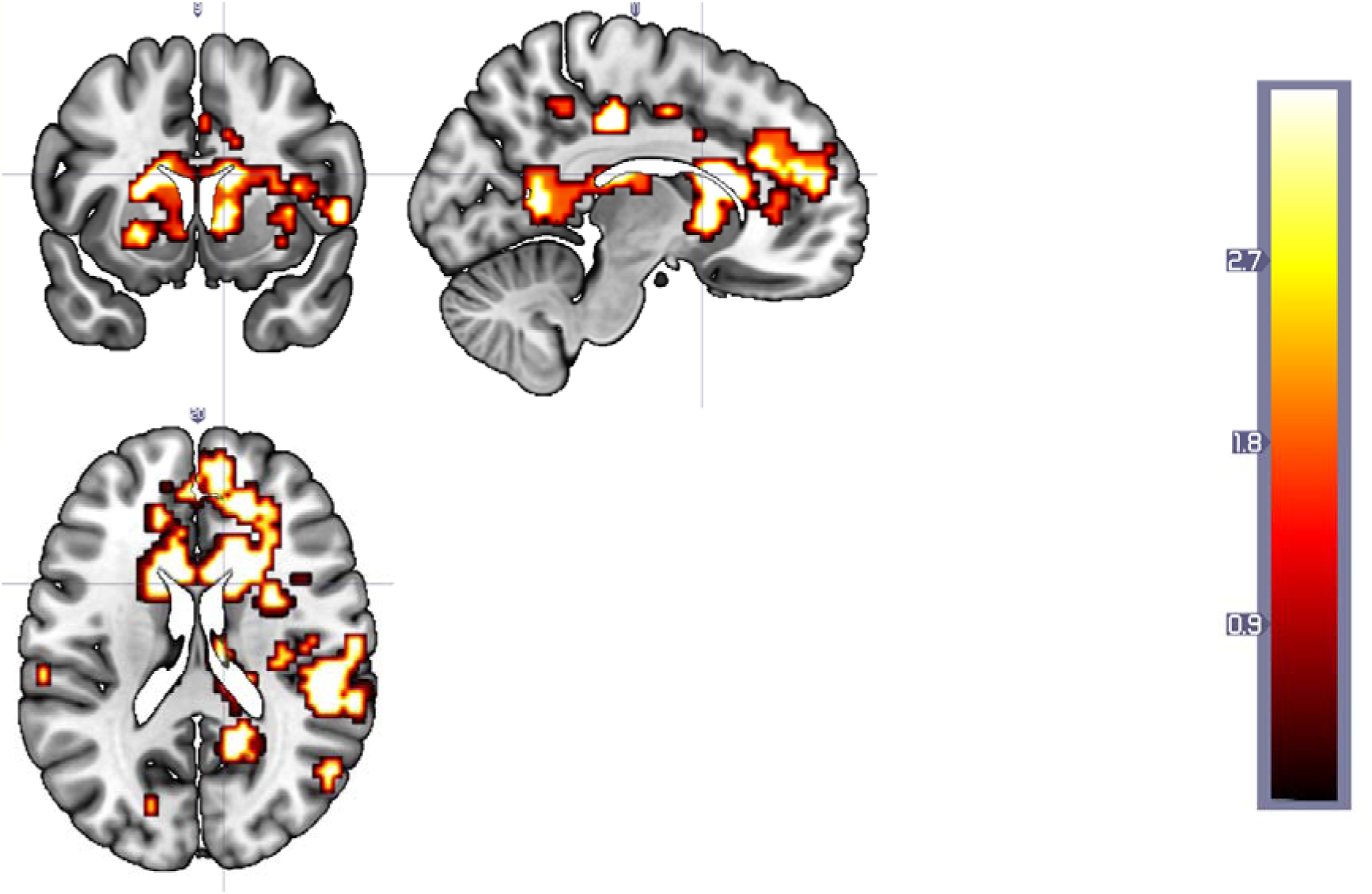
Increased neural activation at T2 (as compared to T1) in BP while processing emotional baby stimuli (as compared to neutral baby stimuli) as compared to CG.

**Table 5.**
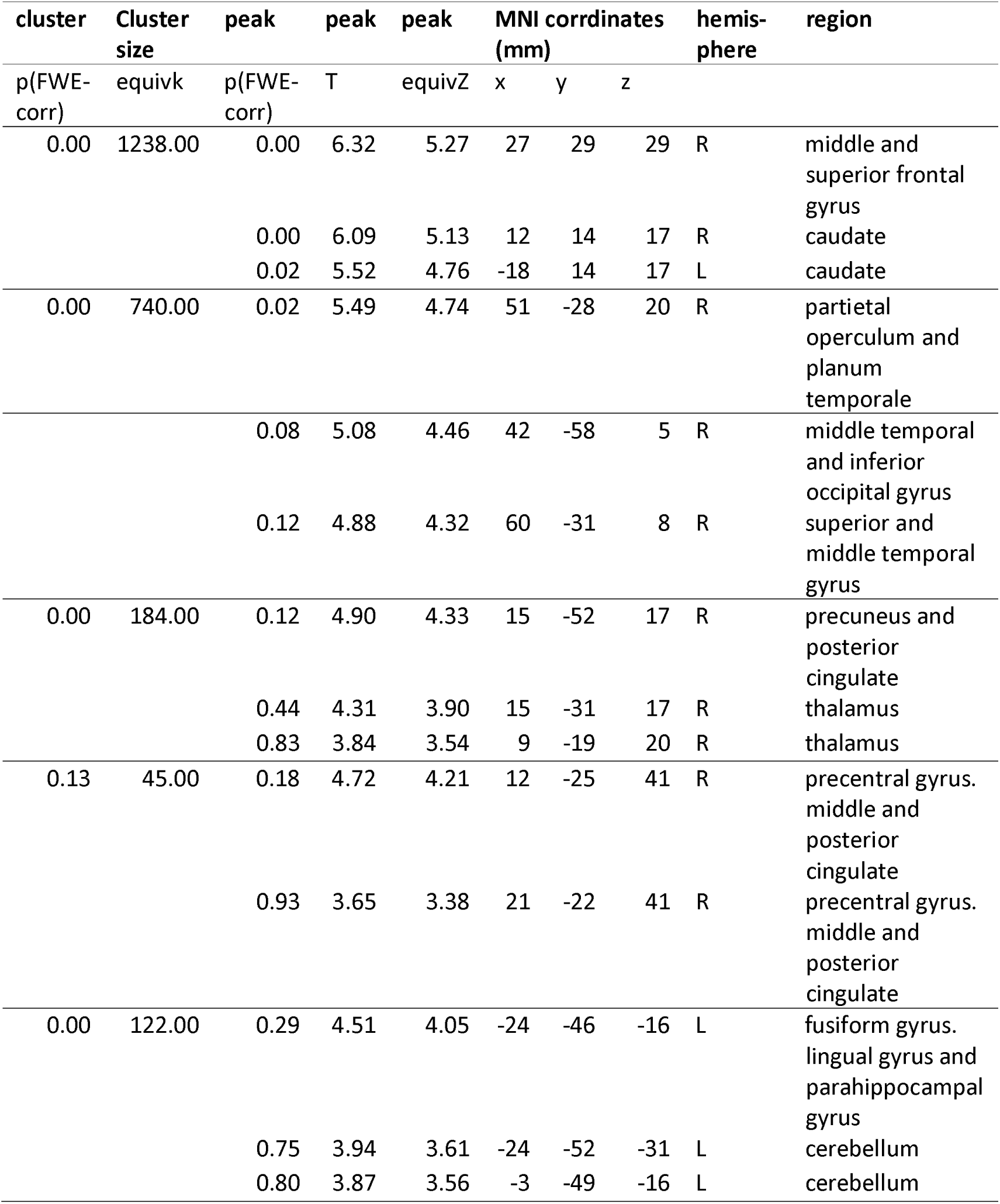
Stronger activation for T2 than T1 for BP larger than CG in the response to emotional baby faces as compared to neutral baby faces.

| cluster | Cluster size | peak p(FWE-corr) | peak T | peak equivZ | MNI coordinates (mm) |  |  | hemisphere | region |
| --- | --- | --- | --- | --- | --- | --- | --- | --- | --- |
| p(FWE-corr) | equivk | p(FWE-corr) | T | equivZ | x | y | z |  |  |
| 0.00 | 1238.00 | 0.00 | 6.32 | 5.27 | 27 | 29 | 29 | R | middle and superior frontal gyrus |
|  |  | 0.00 | 6.09 | 5.13 | 12 | 14 | 17 | R | caudate |
|  |  | 0.02 | 5.52 | 4.76 | -18 | 14 | 17 | L | caudate |
| 0.00 | 740.00 | 0.02 | 5.49 | 4.74 | 51 | -28 | 20 | R | parietal operculum and planum temporale |
|  |  | 0.08 | 5.08 | 4.46 | 42 | -58 | 5 | R | middle temporal and inferior occipital gyrus |
|  |  | 0.12 | 4.88 | 4.32 | 60 | -31 | 8 | R | superior and middle temporal gyrus |
| 0.00 | 184.00 | 0.12 | 4.90 | 4.33 | 15 | -52 | 17 | R | precuneus and posterior cingulate |
|  |  | 0.44 | 4.31 | 3.90 | 15 | -31 | 17 | R | thalamus |
|  |  | 0.83 | 3.84 | 3.54 | 9 | -19 | 20 | R | thalamus |
| 0.13 | 45.00 | 0.18 | 4.72 | 4.21 | 12 | -25 | 41 | R | precentral gyrus. middle and posterior cingulate |
|  |  | 0.93 | 3.65 | 3.38 | 21 | -22 | 41 | R | precentral gyrus. middle and posterior cingulate |
| 0.00 | 122.00 | 0.29 | 4.51 | 4.05 | -24 | -46 | -16 | L | fusiform gyrus. lingual gyrus and parahippocampal gyrus |
|  |  | 0.75 | 3.94 | 3.61 | -24 | -52 | -31 | L | cerebellum |
|  |  | 0.80 | 3.87 | 3.56 | -3 | -49 | -16 | L | cerebellum |

#### 12 months postpartum

Comparing T3 to T1, patients still had increased activation in response to emotional baby faces as compared to controls in regions like temporal gyrus, caudate and posterior cingulate, yet this difference was not as large as at T2, see Figure 9 and Table 6.

**Figure 9.**
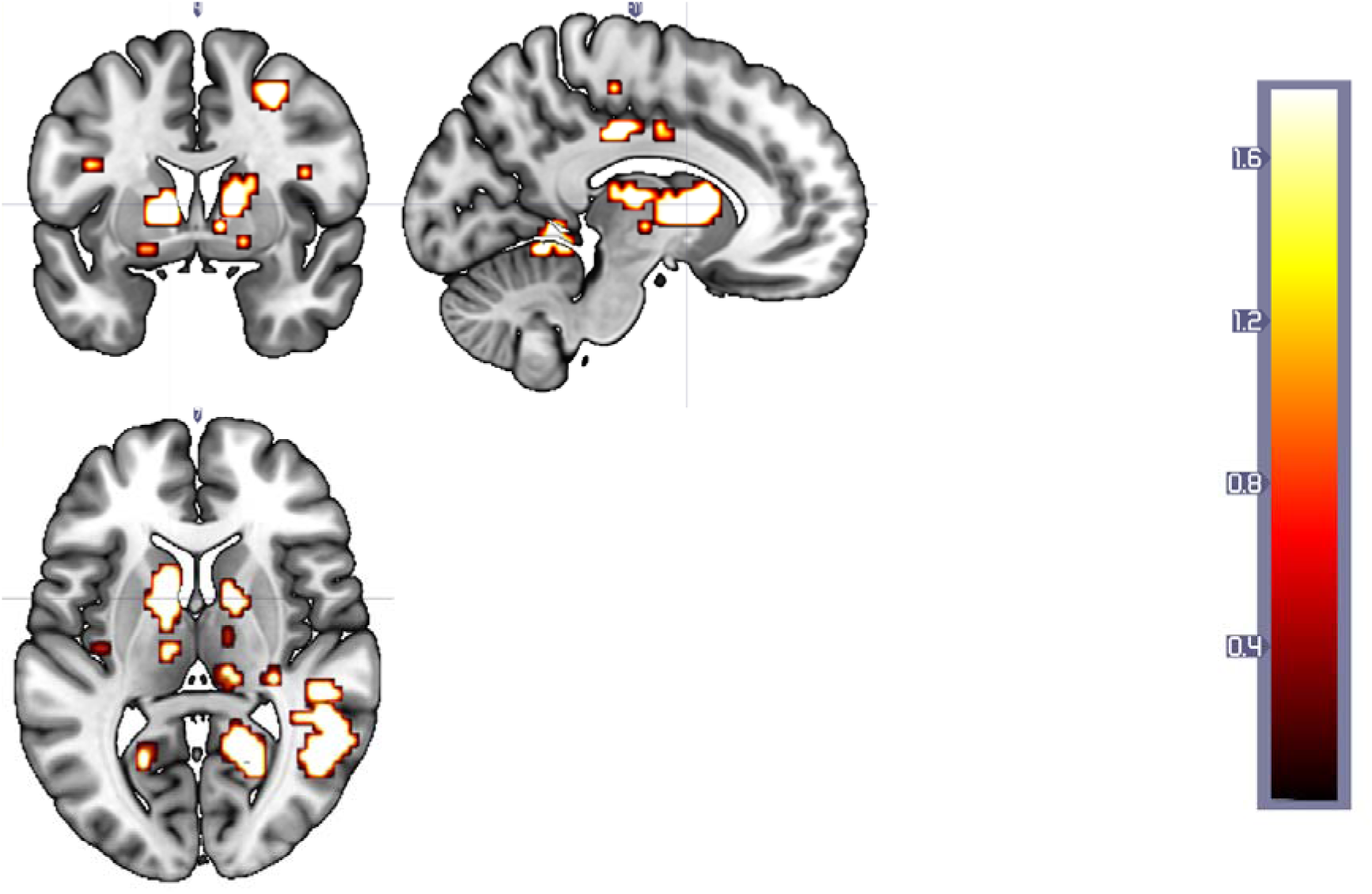
Activation stronger at T3 than T1, larger for the BP mother than for control in response to emotional baby stimuli as compared to neutral stimuli.

**Table 6:** Stronger activation for T3 than T1 for BP larger than CG in the response to emotional baby faces as compared to neutral baby faces.

| cluster | cluster | peak | peak | peak |  |  |  |  |  |
| --- | --- | --- | --- | --- | --- | --- | --- | --- | --- |
| p(FWE-corr) | equivk | p(FWE-corr) | T | equivZ | x.y.z<br>mm | x.y.z<br>mm | x.y.z<br>mm |  |  |
| 0.00 | 340.00 | 0.09 | 4.98 | 4.39 | 57 | -46 | 14 | R | middle and superior temporal gyrus and angular gyrus |
|  |  | 0.14 | 4.80 | 4.26 | 63 | -37 | 14 | R | superior and middle temporal gyrus and supramarginal gyurs |
|  |  | 0.34 | 4.40 | 3.97 | 48 | -43 | 23 | R | angular, superior temporal and supramarginal gyurs |
| 0.34 | 28.00 | 0.11 | 4.88 | 4.32 | 42 | -40 | -13 | R | fusiform and inferior temporal gyrus |
| 0.03 | 82.00 | 0.24 | 4.56 | 4.08 | -12 | 8 | 8 | L | caudate and putamen |
|  |  | 0.40 | 4.32 | 3.91 | -12 | -1 | 5 | L | thalamus, pallidum and caudate |
| 0.00 | 179.00 | 0.29 | 4.47 | 4.02 | 24 | -61 | 8 | R | calcarine cortex, precuneus and lingual gyrus |
|  |  | 0.42 | 4.29 | 3.89 | 12 | -52 | -1 | R | lingual gyrus and posterior cingulate |
|  |  | 0.67 | 4.00 | 3.66 | 15 | -52 | 14 | R | precuneus and posterior cingulate gyrus |

No significant group effects for Go vs. NoGo or the valence of the facial expression was found. Using EPDS as covariate did not result in other or EPDS-related effects for any of the comparisons.

## Discussion

In this report, we present and validate an adapted version of the GoNoGo Task including infant stimuli in a sample mothers with bonding problems (BP) towards their infant. The Infant Emotional GoNoGo Task tests specifically for the perception of and reaction to infant faces with positive, negative, and neutral valence. Our general results suggest that inhibition (NoGo condition) to infant faces activates a broad network for emotion processing both in mothers with and without bonding disorder, including the frontal gyrus, putamen, insula and cingulate cortex. In contrast to the adult faces condition, this activity was even stronger in superior frontal gyrus, nucleus caudatus, accumbens and putamen during the infant faces. This neural network is in line with similar studies using infant stimuli (e.g. (32)), and indicates a high salience of emotional infant faces in particular in mothers in the postpartum period. On a behavioral main effect level, mothers showed significantly higher error rates for baby faces compared to adult faces in the emotional NoGo condition, indicating a higher effort in emotional inhibition for baby faces.

Regarding group differences at 3 months postpartum, on a behavioral level, mothers with BP showed longer reaction times for positive emotional adult faces irrespective of current depressive symptoms as compared to CG, indicating a deficit in the processing of positive emotional stimuli in mothers with BP. Additionally, in the emotional NoGo condition, the inhibition towards emotional baby stimuli was impaired in BP as evident in more frequent mistakes compared to healthy participants. This could be interpreted as an even higher effort to process and react to baby faces in mothers with BP as compared to healthy mothers, which aligns with similar research. At 6 and 12 months, mothers with BD and healthy mothers did not differ regarding reaction times or error rates for emotional adult or baby faces. The immediate reaction to emotional stimuli is not predetermined by the sensation of the stimulus but can be modulated by person factors such as psychopathology and current context under the influence of the central limbic system (59). Both the processing and reaction to emotional stimuli are however a relevant precursors of broader emotion regulation capacities (60). Challenges in emotion regulation in general and specifically the perception of positive affect in others are reported for clinical populations, but the specific behavioral patterns have not yet been shown for BP.

On a neural level, similar to the behavioural data, a difference 3 months postpartum between groups shows that neural response networks comprising insula, cingulate cortex and caudate are stronger activated towards emotional infant stimuli in BP than in CG. While the anterior insular cortex is prominently known for its role in social emotions, especially in anticipating and learning from others’ mental states, the ACC is considered a central node of the emotional processing and regulation network to control affective states on a limbic system level and the caudate as part of the dorsal striatum is considered part of the dopaminergic motivational network. All of them are not only functionally connected but also the network has been discussed to be altered in psychopathology. In addition, they are also involved in the parental brain network that is supposed to be strengthened after becoming a mother while depressed mothers show hypoactivation in the associated brain regions. The evidence of stronger activations of this network could therefore be a correlate of higher effort and usage of neural resources in order to cope with the affective content and is in contrast to the hypoactivation reported for mothers with postpartum depression, thus indicating different mechanisms underlying postpartum depression and BP.

Towards 6 months postpartum, the difference between BP and CG in brain activity has even increased in the aforementioned areas, possibly following the neurofeedback intervention and therefore to be interpreted with caution. Together with the behavioral adjustment, this might be an effect of strong compensation and experience-based improvement in infant emotion processing during the intense first months of motherhood. The training in the daily life of mothers with BP might have stimulated the usage of their neural systems (19). It has been a more general observation for persons with psychological disorders to need increased brain activity for some disorder-specific tasks (for instance for self-referential processing in social anxiety, or working memory in obsessive-compulsive disorder). Towards 12 months postpartum, the neural activation patterns of the BP participants came again closer to the neural pattern of the CG, leaving still group differences in e.g. cingulate and caudate, yet less pronounced than at 6 months. As no intervention took place between 6 and 12 months, this might be interpreted as a normalization of neural systems as parallelly the maternal bonding is less problematic than at previous months (as self-reported in interview based on Brockington/PBQ) and the adaptation to new situation has been mostly successful (see other publications from our team).

Importantly, all behavioural and neural group differences in our data were irrespective of depressive symptoms, although the sample had subclinical depressive scores (that were close to the cut-off score of the EPDS). This adds to the assumption that bonding disorder is clearly distinct from postpartum depression and needs to be considered as own specific clinical field (48).

Taken together we are able to show that neural activation associated with emotion processing baby stimuli is stronger and behavioral performance is impaired in mothers with BP as compared to healthy controls 3 months postpartum and that these findings are independent of depressive symptomatology in these mothers. During the first year postpartum, behavioral performance assimilates in the group of mothers with BD as compared to healthy mothers, while the difference in neural activation in regions associated with emotion processing first increases and then decreases again at the end of the first year after birth of their child.

Our study has some limitations that need to be taken into account for the interpretation of the data and for the planning of future studies. First, and most importantly, the results reported for the differences between 3 months and 6 months postpartum are confounded with the participation of the mothers with BP in a neurofeedback intervention trial. The two tested interventions in the BP group did not result in any behavioral or neural differences for the present analyses, therefore we do not assume they specifically influenced the task performance or neural networks. However, we cannot rule out any effects of the participation on the 6 months assessment. Another limitation that might hamper generalization of results is the sample characteristics and size. As a criterion for participation, the mothers were burdened on a medium level: Severe cases had to be forwarded to immediate therapeutic treatment, while unclear and very mild reports of problems did not lead to inclusion in the BP group. The broad range of clinical comorbidity, however, did not apparently reduce the effect and might be argued rather as strength of the study and for the robustness of findings. Lastly, the generalization of results is clearly limited to women. Future studies might include fathers and other attachment persons. In addition, assessing emotion processing and brain networks already before birth or even pregnancy might help to understand the specific parental brain adaptations and their causes better, both for healthy and clinical individuals.

Our results have some broader clinical implications as well. Our data indicates deficits of the neural emotion processing network that are specific for infant stimuli and for clinical bonding problems, that might be under the influence of learning/experience-based adaptations during the 1^st^ year postpartum, irrespective of depressive symptoms. This adds to the assumption that BP should not be treated as equivalent to depressivity, but has distinct features - as for instance in emotion processing. The reaction to and regulation of children’s affect might furthermore be a relevant target not only for future research but also for therapeutic interventions in the postpartum period.

## Acknowledgements

We gratefully received funding from the Dietmar-Hopp-foundation (ALZ, BD, ME), the German Academic Exchange Service (DAAD, ME), the Heidelberg University mobility program (ME). None of these funding agencies had an influence on recruitment, conducting the study or interpretation of the results.

For their support in study organization, we want to thank our team members Britta Zipser, Nora Nonnenmacher, Leslie Clair, Antonia Huge, Johanna Jübner, Marvin Gänsmantel, Josephine Patrol, Hannah-Sophie Lässig, Franziska Ritter, Julius Zimmer, Linda Stürmlinger and Franziska Schommer

## Author contributions

ME, ALZ and BD designed the study. ME programmed the task. ME, MK, and IB collected the data/did the assessments. ME and MK analyzed the data. ME, MK, BD and ALZ interpreted the results. ME, IB, MK and ALZ drafted the manuscript. All authors contributed and reviewed the final version of the manuscript.

